# Does variation in glucocorticoid regulation predict fitness? A phylogenetic meta-analysis

**DOI:** 10.1101/649608

**Authors:** Laura A. Schoenle, Cedric Zimmer, Eliot T. Miller, Maren N. Vitousek

## Abstract

Glucocorticoid hormones (GCs) are central mediators of metabolism and the response to challenges. Because circulating levels of GCs increase in response to challenges, within-population variation in GCs could reflect individual variation in condition or experience. At the same time, individual variation in the degree to which GCs increase in response to challenges (which is relatively consistent within individuals over time) could have causal effects on stress coping capacity, and the ability to survive and reproduce. Although a number of studies in vertebrates have tested whether within-population variation in GCs predicts components of fitness, it is not clear whether there are consistent patterns across taxa. Here we present the first phylogenetic meta-analysis testing whether within-population variation in GCs is associated with components of fitness across vertebrates. At the same time, we introduce and test predictions about an overlooked but potentially important mediator of GC-fitness relationships: life history context. We suggest that strong context-dependence in the fitness benefit of maintaining elevated GCs could obscure consistent patterns between GCs and fitness across taxa. Meta-analyses revealed that across vertebrates, baseline and stress-induced GCs were consistently negatively correlated with reproductive success. This relationship did not differ depending on life history context. In contrast, the relationships between GCs and survival were highly context dependent, differing according to life history strategy. Both baseline and stress-induced GCs were more strongly negatively associated with survival in longer-lived populations and species. Stress-induced GCs were also more negatively associated with survival in organisms that engage in relatively more valuable reproductive attempts. Fecal GCs did not predict survival or reproductive success. We also used a meta-analytic approach to test whether experimental increases in GCs had consistent causal effects on fitness. Experimental increases in GCs reduced both survival and reproductive success, although the latter relationship was not significant when accounting for phylogeny. Overall, these results support the prediction that GC-fitness relationships can be strongly context dependent, and suggest that incorporating life history may be particularly important for understanding GC-survival relationships. Future studies that explore the role of other aspects of context (e.g., the nature and frequency of stressors, environmental variation) within and across species could provide important insights how and when variation in GC regulation predicts fitness.

## I. INTRODUCTION

In vertebrates, glucocorticoid (GC) hormones play a critical role in mediating the phenotypic plasticity required to live in a dynamic environment. When an individual detects a challenge in the environment, corresponding changes in GCs enable the coordination of a suite of physiological and behavioral traits to match current conditions. GCs support responses to energetically expensive challenges that are predictable (e.g., feeding offspring: (Bonier *et al*., 2009b)) or unpredictable in nature (e.g., winter storms or predation risk: (Boonstra, 2012; Ramenofsky & Wingfield, 2017)). GCs also coordinate life history trade-offs and transitions among life history stages (Wada, 2008; Crespi *et al*., 2013). These presumably adaptive functions of GCs are highly conserved across vertebrates, but maintaining high GC concentrations in order to cope with challenges can also impose costs. GCs can accelerate senescence, increase disease risk, and damage tissues, such as the brain, that are integral to organismal function (McEwen, 2008; Haussmann & Marchetto, 2010; Angelier *et al*., 2018). As a result, the way individuals regulate GC concentrations influences not only their ability to navigate challenges in the short term, but also fitness.

What is the optimal GC-response to challenges? How should GCs be regulated to best match phenotype to the environment while also minimizing the costs of elevated GC concentrations? Answering these questions requires understanding how GC regulation relates to two components of fitness: survival and reproductive success. Describing these relationships has proved challenging in part because of differences in the way GCs shape different components of fitness. A recent meta-analysis in seabirds demonstrated that the relationship between baseline GCs measured in plasma and fitness varied widely depending on the specific metric used to estimate fitness (Sorenson *et al*., 2017). Overall, baseline GCs were negatively associated with reproductive success and food availability, but were not associated with other measures often thought to be indicative of survival ability, such as body condition and foraging effort or reproductive success, including parental care behaviors (Sorenson *et al*., 2017). When considering the numerous empirical studies addressing these relationships using similar fitness measures, variation in environmental conditions as well as individuals’ physiological conditions, life history strategy and evolutionary history have made it challenging to draw generalizable conclusions. Previous qualitative reviews of GC-fitness relationships (Breuner, Patterson, & Hahn, 2008; Bonier *et al*., 2009a; Crespi *et al*., 2013; Schoenle, Zimmer, & Vitousek, 2018a) suggest that the relationships between GCs and both survival and reproductive success are not consistent and vary across these contexts. However, we know of no comparative studies to date that have investigated how context affects GC-fitness relationships.

Here we use a phylogenetic meta-analysis approach to test: 1) whether GC levels are consistently related to reproductive success and survival across vertebrates, and 2) whether these GC-fitness relationships vary with life history traits. We first test the relationships between natural variation in baseline, stress-induced, and fecal measures of GCs and fitness metrics. Next, we determine whether GC manipulations have consistent impacts on survival and reproduction across species. Then, we use species- and population-level life history data to test for a role of life history in influencing GC-fitness relationships. We identify how GC-fitness relationships vary with three components of life history: sex, longevity, and the value of a reproductive bout. A large number of studies addressed the relationship between natural variation in baseline GCs and reproductive success among avian species (Table 1), and thus, we also evaluate how GC-fitness relationships differ across two life history sub-stages of breeding birds: incubation of eggs and chick-rearing.

**Table 1.** The distribution of effect sizes from observational studies included the meta-analyses by taxa, fitness measure, and GC sample type.

| Taxa | Fitness Measure |  |  |  | Sample Type |  | Life History Stage of Sample Collection |  |  |  |
| --- | --- | --- | --- | --- | --- | --- | --- | --- | --- | --- |
|  | Total Effect Sizes | Species | Survival | Reproduction | Baseline Plasma | Stress-Induced Plasma | Fecal | Breeding | Non-breeding | Year-round |
| Birds | 88 | 24 | 22 | 66 | 60 | 25 | 3 | 84 | 4 | 0 |
| Mammals | 19 | 7 | 10 | 9 | 0 | 0 | 19 | 15 | 2 | 2 |
| Reptiles | 3 | 2 | 2 | 1 | 2 | 1 | 0 | 3 | 0 | 0 |
| Fish | 6 | 3 | 1 | 5 | 5 | 1 | 0 | 6 | 0 | 0 |
| Total | 116 | 36 | 35 | 81 | 67 | 27 | 22 | 108 | 6 | 2 |

### (1) GCs and fitness

The mechanisms of GC action are important to understanding the relationships between GCs and fitness in vertebrates. At baseline levels, GCs reflect the overall energetic demands facing an animal, fluctuate with daily and seasonal rhythms, and change across life history stages (Landys, Ramenofsky, & Wingfield, 2006). When an individual perceives a challenge, the hypothalamic-pituitary-adrenal/interrenal (HPA/HPI) axis initiates a hormonal cascade that leads to the secretion of GCs. If the challenge is an immediate threat – such as a predator attack – GC concentrations increase rapidly, and are subsequently downregulated via negative feedback (Sapolsky, Romero, & Munck, 2000; Myers, Mcklveen, & Herman, 2012; Cockrem, 2013). GC regulation is often assessed by measuring plasma GC levels as either baseline (circulating concentrations in the absence of an acute challenge) or stress-induced (circulating concentrations after a standardized restraint stress). GC levels can also be measured in other tissues or matrices, including fecal and feather GCs (which are assumed to reflect GCs over longer time periods).

Baseline GC concentrations tend to mediate aspects of behavior and physiology related to maintaining energetic balance, including foraging behavior, body mass, and metabolism. The link between GCs and energetic challenges led to the hypothesis that individuals with higher baseline GCs are facing more challenging conditions, and thus, will have lower reproductive success and survival rates (reviewed in Bonier *et al*., 2009a). However, because GCs mediate behaviors and physiological functions that can help individuals overcome challenges, higher baseline GCs could also increase reproductive success or survival rates depending on the nature of the challenge(s) individuals face (Bonier *et al*., 2009a; Schoenle *et al*., 2018). For example, in some avian species, higher baseline GCs predict greater reproductive success during the energetically expensive life history stage in which parents are feeding their offspring, presumably because GCs help mediate the physiological and behavioral phenotypes needed to care for young (Bonier *et al*., 2009b; Burtka, Lovern, & Grindstaff, 2016; Henderson *et al*., 2017).

Stress-induced GC concentrations have long been thought to mediate the central life history trade-off between survival and reproduction in favor of immediate survival (Wingfield & Romero, 2001). At the time of the last comprehensive review (Breuner *et al*., 2008) there were few empirical tests of the relationship between stress-induced plasma GCs and either survival or reproductive success. The prevailing theory has remained that stress-induced GCs promote survival to the detriment of non-essential activities such as reproduction (Schoenle *et al*., 2018). However, whether or not stress-induced GCs will promote survival could depend on the nature of the challenge(s) and the frequency or intensity of the challenge(s) (Schoenle *et al*., 2018). GC-mediated changes in phenotype will not improve chances of survival if the challenge cannot be escaped, avoided, or overcome. For example, if individuals are at risk for starvation and no food is available, the challenge cannot be escaped, and higher GCs will be associated with starvation risk, and likely mortality (Wingfield *et al*., 1998; Romero & Wikelski, 2010). Furthermore, evidence suggests that high GC reactivity in response to stressors generally promotes survival in high risk environments, but might prove costly in low risk environments (Schoenle *et al*., 2018). For example, among American redstarts (*Setophaga ruticilla*) overwintering in low-quality environments, those with higher stress-induced GC levels are more likely to survive; however, there is no relationship between stress-induced GCs and survival among birds overwintering in environments with higher food availability (Angelier, Holberton, & Marra, 2009).

As stress-induced GCs are assumed to promote survival to the detriment of reproduction, high stress-induced GC levels are usually associated with reduced reproductive success. For example, individuals that respond to a standardized stressor with higher stress-induced GC levels have lower fledging success in female barn swallows (*Hirundo rustica erythrogaster*) (Vitousek, Jenkins, & Safran, 2014), and higher rates of nest desertion in male great tits (*Parus major*) (Ouyang, Quetting, & Hau, 2012). Therefore, it has been proposed that breeding individuals should reduce their acute stress response when the value of the current reproductive bout is high (Bókony *et al*., 2009). Lapland longspurs (*Calcarius lapponicus*) attenuate their stress response when facing a snow storm during the parental stage but not during the pre-parental stage (Krause *et al*., 2016). However, if mounting a strong stress response during breeding allows individuals to cope efficiently with challenges and then resume breeding activities, high stress-induced GCs can potentially be positively associated with reproductive success. For example, female tree swallows (*Tachycineta bicolor*) that show a strong stress response to a standardized stressor are less likely to abandon their nest during incubation under challenging conditions – as long as this response is coupled with the ability to effectively terminate the stress response through negative feedback (Zimmer *et al*., 2019).

Because the benefit of elevated GCs is expected to vary across contexts, and life history can influence GC-fitness relationships (described below), we predict that meta-analyses will show context-dependent rather than consistent relationships between baseline GCs or fecal GCs and metrics of fitness. Similarly, we predict that analyses that do not incorporate context dependence will find no consistent relationship between variation in stress-induced GCs and survival; however, we predict that individuals that maintain higher stress-induced GCs during reproduction will have consistently lower reproductive success.

### (2) Context-dependence in GC-fitness relationships: a role for life history

The fields of evolutionary and behavioral endocrinology have long posited that the relationships between hormones and fitness depend on context, and in particular, life history (Breuner, 2010; Williams, 2012). However, few studies have empirically tested how GC-fitness relationships vary with life history traits. Among those that have, the effects of GCs on survival and reproduction differ with life history strategy. For example, in both side blotched lizards (*Uta stansburiana*) and black-legged kittiwakes (*Rissa tridactyla*), experimental increases in GCs reduce reproduction in groups of animals (color morphs in the lizards and populations in kittiwakes) demonstrating slower paces-of-life (i.e., more K-selected strategies), but not in individuals with faster paces-of-life (i.e., r-selected strategies) (Lancaster *et al*., 2008; Schultner *et al*., 2013). Large-scale, cross-species analyses will be necessary to understand if these patterns are generalizable. Here, we test whether life history traits – including longevity and the value of a single reproductive bout – influence GC-fitness relationships across species. We include sex as a factor in our analyses, but we do not make explicit predictions for how sex should shape the GC-fitness relationship, as we predict this will be highly dependent on the breeding system.

Longevity could shape the relationship between GC concentrations and fitness in several ways. Because GCs are thought to promote survival in the face of challenges (Sapolsky *et al*., 2000; Breuner *et al*., 2008), we might expect selection to favor individuals with elevated GC levels. If so, longer-lived species could have elevated GC concentrations (Hau *et al*., 2010), and potentially, more positive GC-survival relationships within populations. Alternatively, maintaining high concentrations of glucocorticoids could impose physiological costs that accumulate over time (McEwen & Seeman, 2004), and thus longer-lived animals could suffer greater costs of high GCs than shorter-lived animals (Schoenle *et al*., 2018). If GCs have cumulative costs, we predict that longer-lived animals will have more negative GC-fitness relationships, both in terms of survival and reproduction. While a recent phylogenetic comparative analysis tested this prediction across species (Vitousek *et al*., 2019), we know of no studies that have investigated how the relationships between individual variation in GCs and fitness metrics vary with longevity across species.

The value of a reproductive bout is also a critical life history trait that is connected to pace-of-life. Species with faster paces-of-life tend to have fewer reproductive bouts and produce more offspring each bout than those with slower paces-of-life; thus, on average each reproductive attempt represents a greater proportion of total lifetime reproductive effort in these species (higher reproductive value: (Bókony *et al*., 2009; Vitousek *et al*., 2019)). Because acutely elevated GCs often downregulate reproduction in favor of survival, we generally predict a more negative relationship between stress-induced GCs and reproductive success when reproductive value is high. However, in species that engage in particularly high value reproductive attempts, selection could also favor individuals in which acutely elevated GCs do not downregulate reproduction (Boonstra & Boag, 1992; McConnachie *et al*., 2012). In these species, we would expect no relationship between stress-induced GCs and reproductive success. Because elevated baseline GCs may help to support periods of energetic challenge, we predict more positive baseline GC-reproductive success relationships in species with higher reproductive value (Bókony *et al*., 2009; Schoenle *et al*., 2017). We do not expect that relationships between baseline GCs and survival will differ according to reproductive value.

Because of the role of baseline GCs in supporting energetically demanding periods, we also predict that GC-fitness relationships will differ across life history stages (Bokony *et al*. 2009). In birds, we predict less negative relationships between baseline GCs and reproductive success during the offspring provisioning period than during incubation, because elevated baseline GCs could help to support this energetically demanding period (Bonier *et al*., 2009b).

### (3) Hormone Manipulations and Fitness

In addition to measuring correlations between natural variation in GCs and fitness measures, some studies have manipulated GCs to determine whether or not GCs can cause increases or decreases in survival or reproductive success. Ideally, GC manipulations would confirm the results from studies of natural GC variation, and predictions for the relationships between experimental elevations of GCs and fitness metrics would mirror those for natural variation in GCs. However, hormone manipulations might not simply produce the same results as natural variations in concentrations of GCs. Nearly all GC-manipulation studies in wildlife use techniques such as slow-release implants, topical treatments, food supplementation, or injection to increase GC concentrations (Sopinka *et al*., 2015). If GC manipulations can successfully elevate GCs within the normal physiological range for a species – a common concern for such studies – most manipulations remain unable to replicate the pulsatile, cyclic release of endogenous hormones (Fusani, 2008; Dantzer, Westrick, & van Kesteren, 2016). Therefore, experimental manipulations of GCs could influence not just hormone concentrations, but the entire HPA axis, including receptor expression and negative feedback, thus altering GC’s downstream effects (Fusani, 2008). Furthermore, hormone regulation has been selected over evolutionary time, and as a result, GC manipulations that move individuals’ hormone concentrations away from their natural concentrations could have negative effects on multiple aspects of fitness (Crossin *et al*., 2016; Schoenle *et al*., 2019). Thus, we predict that experimental increases in GCs will reduce both survival and reproduction.

### (4) GC-fitness meta-analyses

To evaluate how GCs relate to reproductive success and survival across vertebrates we performed eight meta-analyses of the following relationships: 1) baseline GCs and survival, 2) baseline GCs and reproduction, 3) stress-induced GCs and survival, 4) stress-induced GCs and reproduction, 5) fecal GC metabolites and survival, 6) fecal GC metabolites and reproduction, 7) experimentally manipulated GCs and survival, and 8) experimentally manipulated GCs and reproduction. To test whether GC-fitness relationships vary with context, specifically life history, we used moderator analyses to test how longevity, the value of a reproductive bout, and sex relate to relationships between GCs and both survival and reproductive success. We were further able to test predictions about the role of life history substages in influencing baseline GC-reproductive success relationships using the large number of effect sizes present in the avian literature (*n* = 55). Here we conducted a meta-analysis of baseline GC-reproductive success within birds alone that included an additional moderator: breeding life history substage (incubation or chick-rearing). Finally, we tested for an effect of publication bias on the results of each analysis.

## II. METHODS

### (1) Literature Search

We performed a systematic review of the literature assessing the relationship between GCs and fitness metrics including both observational studies and hormone manipulation experiments. We identified relevant research by searching Web of Science using combinations of the terms “gluc*,” “cort*,” “fitness,” “survival,” and “reproduction,” and searching Google Scholar (which does not allow for searching with truncations and wildcards), using many variants of the hormone names (e.g., GC, GCs, glucocorticoids, cortisol). Additionally, we searched forward and reverse citations of review and synthesis papers addressing the GC-fitness relationship including: Breuner *et al*. 2008, Bonier *et al*. 2009, Hau *et al*. 2010, Sorenson *et al*. 2017. The search included literature published (including online publication) before January 2018.

### (2) Inclusion/Exclusion Criteria

We included studies that met the following inclusion/exclusion criteria: 1) Studies must have been conducted in free living vertebrates. 2) Observational studies must have measured both a fitness metric(s) and GC hormones as either baseline plasma GC concentrations, stress-induced plasma GC concentrations, or fecal GC metabolites. 3) Experimental hormone manipulations must have both directly altered GC concentrations by administering a hormone treatment with the goal of increasing GC levels and also assessed fitness metrics post-treatment. Studies that indirectly manipulated GCs (e.g., by food supplementation) were excluded. 4) Studies assessing survival rates must have evaluated mortality measuring return rates, re-sightings, or comprehensive tracking that allowed for determination of death. 5) Studies assessing reproductive success must have measured both GCs and reproductive success within the same reproductive season (i.e., not carry-over effects). 6) Measures of reproductive success must either have been the number of offspring produced or a binary, definitive evaluation of success or failure (e.g., abandoned or successfully reared offspring).

### (3) Data extraction and effect size calculations

We collected effect sizes for the relationship between GCs and fitness metrics from reported test statistics (e.g., *F, χ*^2^, *z, R*^2^, *r*). We identified whether the fitness metric was indicative of survival or reproductive success and recorded the specific fitness measure used. For studies in which GC concentrations were manipulated, we recorded the length of time from administering the treatment until assessing the fitness metric. We also recorded the sample size, degrees of freedom (when relevant), and direction of the GC-fitness relationship. We assigned a positive value for the direction of the relationship when higher GCs were associated with higher rates of survival or reproduction and a negative value when higher GCs were linked to lower rates of survival or reproduction. For 9 effect sizes in which the GC-fitness relationship was not significant, we were unable to identify the direction of the relationship. We ran all analyses including these studies (meta-analyses for baseline GCs and experimental manipulations of GCs) by assigning all effect sizes a negative value or all a positive value (as in Bentz, Becker, & Navara, 2016). The results of the meta-analyses did not differ qualitatively when these studies were assigned all negative or all positive values. We report the results from all analyses assuming a positive relationship. In cases where the statistics were not reported explicitly (often the case in publications using model selection as the primary analytic tool), but data were available in a repository, we used the raw data to calculate the correlation coefficient *r*. Alternatively, if the data were not available in a repository, but were presented in a scatterplot, we used WebPlotDigitizer version 3.9 (Ankit Rohatgi, Austin, Texas, USA) to extract data and calculate *r*. When statistics were not reported and we were unable to access the data from a repository or graph, we contacted the corresponding author and requested data or statistics (25 effect sizes from 5 studies were obtained from author inquiries). The sources for each effect size are included in the supplementary material.

All effect sizes were converted to the correlation coefficient *r* using formulas from: Rosenthal & Dimatteo (2001), Nakagawa & Cuthill (2007), and Rosenberg, Rothstein, & Gurevitch (2013). There is no widely accepted way to identify degrees of freedom from random effects models. Thus, we followed a technique used in recent meta-analyses (Bentz *et al*., 2016), and when calculating effect sizes from random effects models we converted the *p-value* to a normal deviate Z-score and calculated *r* as *r*= *Z*/√*N* (Rosenthal & Dimatteo, 2001). If the *p-value* was reported as less than a value (e.g. < 0.001), then we used that value to calculate *r*.

To avoid pseudo-replication, each effect size was assigned a reference ID and a group ID to be used as random effects in the analyses. Reference IDs were a unique number associated with each publication such that all effect sizes from a single publication were assigned the same reference ID. Group IDs indicated if the same individuals were used to calculate multiple effect sizes (e.g., the relationship was measured across multiple years or seasons using the same individuals), either within the same publication or across multiple publications.

For observational studies, we recorded whether the hormone measure was baseline, stress-induced, or from fecal metabolites based on the authors’ classification of the measure. Although not included in the analyses, we also recorded the latency from capture until collecting blood to measure baseline GCs and the time after restraint at which stress-induced GCs were measured. Among studies of baseline GCs, most effect sizes were calculated using GCs measured in plasma samples collected within 3 minutes after capturing the animal (*n* = 52). However, several studies (*n* = 7) measured GCs over a longer time window (e.g., up to 5 minutes) but either verified that concentrations did not increase from baseline levels within that selected time period (e.g. Strasser & Heath, 2013)) or collected most samples in less than 3 minutes, but the maximum latency was longer (e.g. Nelson *et al*., 2015). Although they met all other requirements for the analysis, we excluded 2 studies that collected blood samples 10 minutes after capture (Beletsky, Orians, & Wingfield, 1989, 1992) and did not justify that hormone levels could be considered baseline. Latency to sample for baseline GCs was not recorded for 8 effect sizes, but studies reported collecting the samples immediately after capture (e.g. Magee, Neff, & Knapp, 2006) or using non-invasive sampling methods, such as blood-sucking insects (Bauch *et al*., 2016), and these were retained in the analysis. We also recorded the length of the standardized period of restraint used prior to collecting a blood sample for stress-induced GCs. The time between stressor exposure and peak stress-induced GCs varies across species and individuals, and most species peak after 14-60 minutes of restraint (Breuner *et al*., 2008; Satterthwaite *et al*., 2010; Small *et al*., 2017). We chose to accept authors’ selected timeframes as an acceptable measure for stress-induced GCs. Fecal metabolites were considered an integrative measure of hormones and were not associated with a specific time range.

For each effect size, we collected data on life history variables that could influence the direction or strength of the GC-fitness relationship. We recorded species, sex (male, female, or pooled sexes), life history stage in which the sample was collected (breeding, non-breeding, or year-round). We defined breeding as the periods of time involving preparing for breeding (e.g., mate attraction, nest building), mating, and caring for offspring shortly after birth or hatching (e.g. provisioning young in the nest (birds), lactation (mammals)). Any periods outside of this window were classified as non-breeding. We categorized studies collecting data across breeding and non-breeding stages as year-round. All studies of GC-reproductive success relationships measured GCs during breeding. We also recorded the specific sub-stage of breeding (e.g., lactation, spawning, incubation, chick-rearing). Among the included studies, non-breeding periods were distinguished from breeding periods in that they were in a spatially distinct location, did not involve parental care, or were explicitly identified by the authors as non-breeding.

We collected species- and population-level life history data to test how life history shapes GC-fitness relationships. We obtained these life history variables from the extended HormoneBase dataset (Johnson *et al*., 2018) including: maximum longevity, litter or clutch size, and the number of litters/clutches per year. For any species that were not in HormoneBase, we obtained these life history variables from the Amniote (Myhrvold *et al*., 2015) and AnAge (De Magalhaes & Costa, 2009) databases. From these variables, we calculated the expected value of a single reproductive bout (called “brood value” in the avian literature) following the formula in Bókony *et al*. (2009): log_10_(clutch or litter size/[clutch or litter size x bouts per year x maximum lifespan])

### (4) Building the phylogeny

We built a phylogeny to incorporate the evolutionary relationships among the study taxa following an identical pipeline to that described in Johnson *et al*., (2018). In short, we used manual taxonomic reconciliation to match every study species to a tip in a recent, large taxon-specific phylogeny: ray-finned fishes (Rabosky *et al*., 2013), amphibians (Pyron & Wiens, 2011; Eastman, Harmon, & Tank, 2013), mammals, squamates (Pyron, Burbrink, & Wiens, 2013), turtles (Jaffe, Slater, & Alfaro, 2011), and birds (Jetz *et al*., 2012). We then pruned these taxon-specific trees to the manually matched study species, employing tip-row swaps for phylogenetically equivalent species when necessary (Pennell, FitzJohn, & Cornwell, 2016). Finally, we bound these taxon-specific trees into a backbone, vertebrate-scale phylogeny created with the TimeTree of Life (Kumar *et al*., 2017) such that the final tree was ultrametric and contained one tip for every species in the study.

### (5) Meta-analysis

All analyses were performed in R version 3.5.1 (R Core Team, 2018) and meta-analyses were performed using the package *metafor* (Viechtbauer, 2010). Before performing any analyses, we used the *escalc* function in *metafor* to convert the correlation coefficients *r* to the standardized, normally distributed effect size Fisher’s Z and to calculate the sampling variance for each effect size (as described in Holtmann, Lagisz, & Nakagawa, 2016; Taff, Schoenle, & Vitousek, 2018). Results were back-transformed to r to support easier interpretation.

We performed meta-analyses using linear mixed models using the *rma.mv* function in *metafor* to control for the non-independence of effect sizes collected from the same study, species, or individuals as well as non-independence associated with phylogenetic relationships. Thus, we tested for the importance of the random effects in the models following Foo *et al*., (2016). We tested the importance of the reference ID, species, and whenever relevant, group ID using likelihood ratio tests. We separately tested each random variable for significance within an intercept-only meta-analysis. Variables that were found to be significant in these individual analyses were then included in an intercept-only model. We checked which of these random effects remained significant after accounting for the others, and only retained those random effects in the general models. We report the retained effects in Table 3 and the statistics from each test in the electronic supplementary material.

**Table 3.** The sample size, random effects, phylogenetic variance, and I^2^ for each meta-analysis.

| Meta-analysis | n<br>(Effect Sizes) | Random Effects Retained<sup>1</sup> | Variance ($\sigma^2$) Explained by<br>Shared Evolutionary History<sup>2</sup> | $I^2$ (%)<sup>3</sup> |
| --- | --- | --- | --- | --- |
| Baseline GCs & Survival | 14 | None | 0.02 | 30.81 |
| Baseline GCs & Reproduction | 53 | Group ID | 0.000 | 79.83 |
| Baseline GCs & Reproduction<br>(birds only) | 48 | Group ID | 0.000 | 78.91 |
| Stress-induced GCs & Survival | 11 | None | 0.019 | 57.96 |
| Stress-induced & Reproduction | 16 | None | 0.000 | 59.08 |
| Fecal GCs & Survival | 10 | Reference ID | 0.71 | 92.35 |
| Fecal GCs & Reproduction | 12 | Reference ID | 0.000 | 78.92 |
| Experimentally Elevated GCs &<br>Survival | 6 | None | 0.000 | 75.43 |
| Experimentally Elevated GCs &<br>Reproduction | 15 | Reference ID | 0.13 | 71.08 |

<sup>1</sup>Random effects found to be significant using likelihood ratio tests (statistics reported in the supplementary material). For each analysis we tested for the importance of reference ID, group ID, and species. <sup>2</sup>Variance explained by including a phylogenetic covariance matrix as a random effect in the meta-analysis. <sup>3</sup>The percent of variability in effect sizes due to variation in the effects (rather than sampling error) from an intercept-only model with no random effects.

We determined the overall effect sizes for the relationships between glucocorticoids and fitness with restricted maximum likelihood meta-analytic models including the selected random effects using the *rma.mv* function. We used a phylogenetic meta-analysis to account for the expectation of similar trait values among related species by including these relationships in the form of a phylogenetic correlation matrix random effect. For each meta-analysis, we calculated the heterogeneity statistic I^2^ using the *rma.uni* function (reported in Table 3). I^2^ is the percent of variability in the effect sizes (0-100%) that is due to variation in the effects (rather than sampling error) from an intercept only model with no random effects. This variation in the GC-fitness relationship might be accounted for by other moderators (e.g. sex, life history stage, breeding stage, or other factors).

### (6) Moderator Analyses

For the meta-analyses including baseline, stress-induced, and manipulated GCs, we tested three moderators: sex, the value of a reproductive bout, and longevity. We were unable to test for correlations between sex and fecal GC-fitness relationships because of a lack of effect sizes for both males and females. We conducted all moderator analyses using phylogenetically informed meta-analyses with moderators as fixed effects and the previously selected random effects using the *rma.mv* function. Because the sample size varied for different moderator analyses, we performed a separate meta-regression model for each moderator rather than including all moderators in a single analysis.

The large number of effect sizes from birds allowed us to perform additional moderator analyses to further explore the role of context in GC-fitness relationships. We first performed a baseline GC-reproductive success meta-analysis using effect sizes from birds only. Of the 48 effect sizes, 40 were from distinct substages of breeding: incubation and chick-rearing. Thus, following the main meta-analysis, we were able to run moderator analyses specific to birds, including not only sex, value of a reproductive bout, and longevity, but we also tested whether the relationship between GCs and reproductive success varied across life history substages.

We also performed a moderator analysis using the effect sizes from studies manipulating GC concentrations. We tested whether or not the length of time between the hormone manipulation and when the fitness metric was assessed influenced the relationship between GCs and fitness. If the effects of GCs on fitness are limited to the short-term, then as more time passes between the treatment and assessing fitness measures the effect sizes will approach 0.

### (7) Tests for publication bias

We tested each of the eight meta-analyses for evidence of publication bias in several ways. We performed Egger’s regression test (Egger *et al*., 1997) using the overall intercept-only models with the *regtest* function in *metafor*. Egger’s test evaluates the relationship between standardized residuals and study precision. We also conducted a trim-and-fill analysis using the *trimfill* function in metafor to test for asymmetry among effect sizes in a funnel plot. Funnel plot asymmetry can indicate heterogeneity among effect sizes or publication bias in favor of significant results. Finally, we used a linear regression to evaluate the relationship between publication year and effect size for time-lag bias. Evidence of time-lag bias occurs when effect sizes approach zero as time passes because studies with larger effect sizes are published faster or earlier than those with smaller effect sizes.

## III. Results

### (1) GC-fitness dataset

Our dataset included 137 effect sizes from 59 publications (complete dataset provided in the supplementary material). Table 1 illustrates the distribution of effect sizes from observational studies by taxa, fitness measure, GC sample type, and the life history stage when samples were collected. Table 2 shows the distribution of effect sizes from hormone manipulation studies by taxa and fitness measure. All observational and experimental studies addressing GC-reproductive success relationships were conducted during breeding (i.e., blood sample collection and hormone manipulations were conducted during the same breeding season in that reproductive success was measured). The meta-analyses demonstrated that the relationships between GCs and fitness vary with the hormone sample type and the fitness metric used (Figure 1). The final models for the meta-analysis, including the random effects retained, variance explained by shared evolutionary history, and I ^2^ (the percent of variability in effect sizes due to variation in the effects rather than sampling error), are summarized in Table 3. Details for the tests of significance of random effects are included in the supplementary material.

**Table 2.**
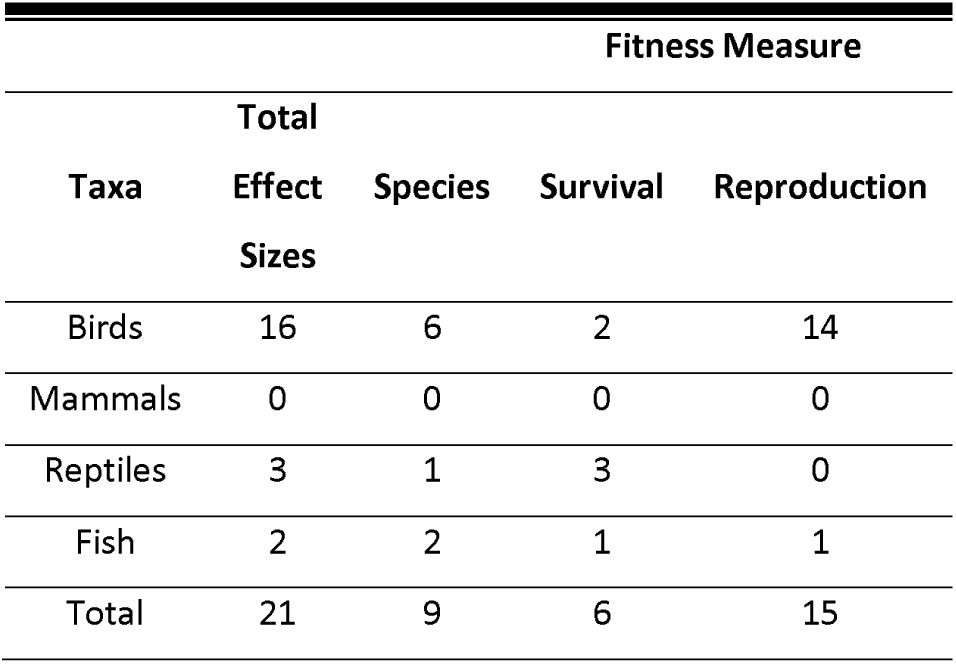
The distribution of effect sizes from hormone manipulation studies by taxa and fitness measure.

**Figure 1.**
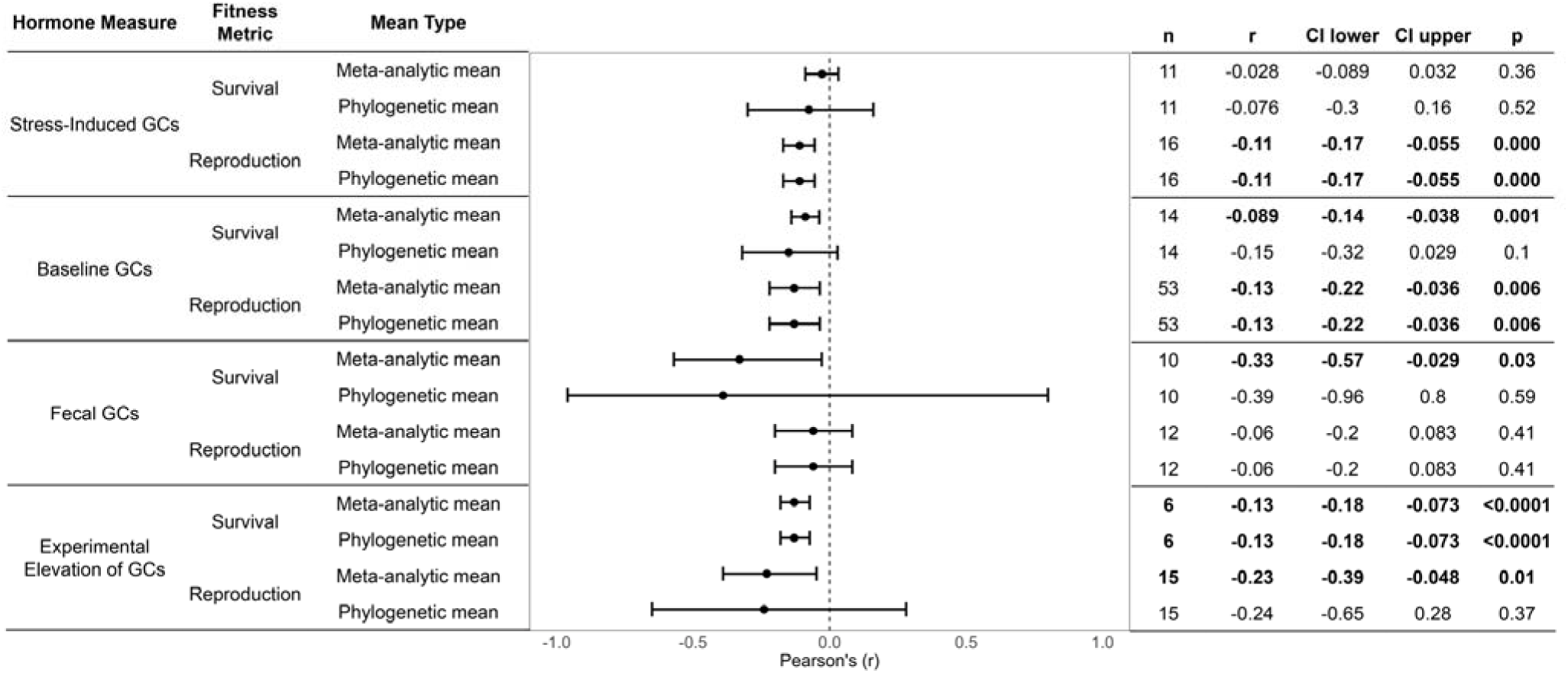
Meta-analytic means and phylogenetic meta-analytic means for studies assessing the relationships between GCs and fitness measures. Means are represented as a point and error bars represent the 95% confidence interval. Significant effects are bolded.

### (2) Baseline GCs and survival

Higher baseline GCs were associated with lower survival, but the effect was small and the relationship was not significant when incorporating the expected covariance between related species due to shared evolutionary history (Figure 1, Table 3). Moderator analyses revealed that populations and species with slower paces-of-life had more negative baseline GC-survival relationships. Longer-lived populations and species had more negative baseline GC-survival relationships than their shorter-lived counterparts (β = −0.01, 95% CI = −0.019 --0.0014, *p* = 0.02, n = 14, Figure 2a). Similarly, animals with a lower value for a single reproductive bout tended to show more negative baseline GC-survival relationships, but the effect was not significant (β = 0.35, 95% CI = −0.02 – 0.64, *p* = 0.06, n = 14, Figure 2b). Sex was not associated with the baseline GC-survival relationship (males relative to females, β = −0.043, 95% CI = −0.21 – 0.12, *p* = 0.61, n = 10).

**Figure 2.**
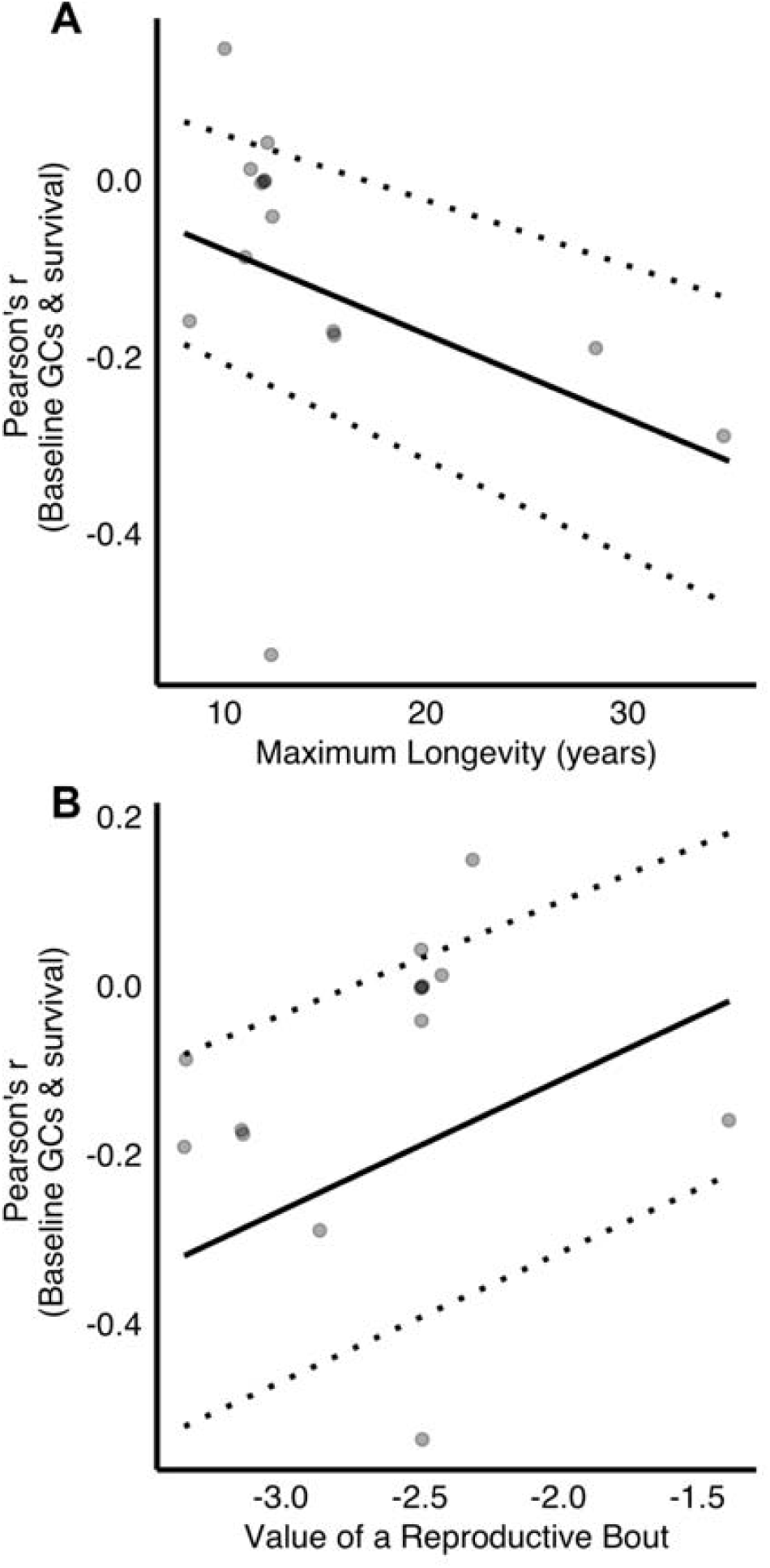
The correlation between longevity (A) and the value of a reproductive bout (higher values indicate a relatively higher value of the current brood) (B) and the relationship between baseline GCs and survival. Each point represents an effect size and the dashed lines indicate 95% confidence intervals.

### (3) Baseline GCs and reproductive success

Overall, higher baseline GC concentrations during reproduction were associated with lower reproductive success; this relationship was identical for the standard and phylogenetically-controlled meta-analyses (Figure 1). None of the moderators accounted for the variation in the relationships between baseline GCs and reproductive success (sex (male): β = 0.076, 95% CI = −0.037 – 0.19, *p* = 0.19, n = 45; reproductive value: β = 0.057, 95% CI = −0.22 – 0.32, *p* = 0.69, n = 53; longevity: β = 0.0012, 95% CI = −0.15 – 0.16, *p* = 0.99, n = 53).

Meta-analyses within birds alone that included breeding life history stage as an additional moderator also revealed a significant negative relationship between baseline GCs and reproduction for both the standard and phylogenetically informed meta-analyses (for both, r = −0.12, 95% CI = −0.21 – 0.029, *p* = 0.01, n = 48). None of the moderators, including breeding life history substage, explained the heterogeneity in GC-reproduction effect sizes (sex: (male)β = −0.079, 95% CI = −0.045 – 0.20, *p* = 0.21, n = 40; breeding substage: (chick-rearing) β = 0.12, 95% CI = −0.0044 – 0.24, *p* = 0.059, n = 40; longevity: β = −0.002, 95% CI = −0.012 – 0.0077, *p* = 0.68, n = 48; value of a reproductive bout: β = −0.05, 95% CI = −0.40 – 0.31, *p* = 0.79, n = 48).

### (4) Stress-induced GCs and survival

Overall, the correlation between stress-induced GCs and survival was not significant. Both the phylogenetically informed meta-analysis and the standard meta-analysis produced qualitatively similar results (Figure 1). Relationships between stress-induced GCs and survival appeared to be strongly context dependent. Studies conducted in longer-lived animals found more negative GC-survival relationships than those conducted in shorter-lived animals (β = - 0.01, 95% CI = −0.02 - −0.002, *p* = 0.01, n = 11, Figure 3a). In addition, animals with a higher value for a single reproductive bout had more positive GC-survival relationships (β = 0.28, 95% CI = 0.06 – 0.48, *p* = 0.01, n = 11, Figure 3b). Sex was not associated with the GC-survival relationship (males relative to females, β = 0.023, 95% CI = −0.12 – 0.16, *p* = 0.74, n = 8).

**Figure 3.**
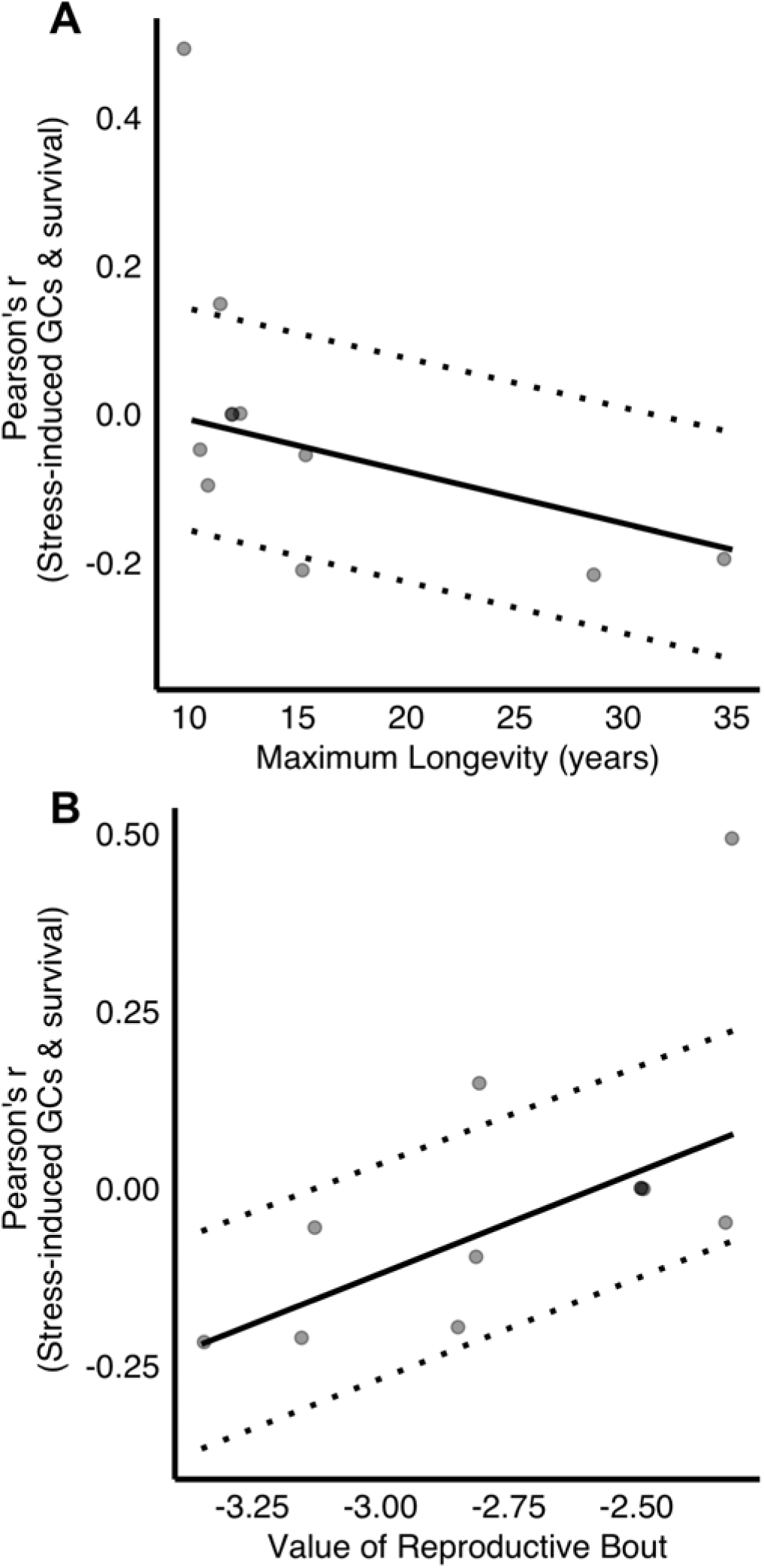
The relationship between stress-induced GCs and survival varies according to (A) longevity and (B) the value of a reproductive bout (higher values indicate a relatively higher value of the current brood). Each point represents an effect size and the dashed lines indicate 95% confidence intervals.

### (5) Stress-induced GCs and reproductive success

Both the phylogenetically informed and standard meta-analyses demonstrate a significant, negative relationship between stress-induced GCs and reproductive success (Figure 1). Moderators did not explain any of the observed variation in effect sizes (sex (male): β = - 0.086, 95% CI = −0.20 – 0.036, *p* = 0.17, n = 15; value of a reproductive bout: β = −0.014, 95% CI = −0.96 – 0.068, *p* = 0.74, n = 16; longevity: β = 0.0062, 95% CI = −0.0085 – 0.021, *p* = 0.41, n = 16).

### (6) Fecal GCs and fitness

Overall, fecal glucocorticoids were negatively associated with survival, but the relationship was not significant after incorporating the expected covariance in traits among related species (Figure 1). Because there were no data for females, we were unable to test for an effect of sex. None of the other moderators (value of a reproductive bout: β = 0.30, 95% CI = −0.57 – 0.85, *p* = 0.53, n = 10; longevity: β = −0.015, 95% CI = −0.05 – 0.015, *p* = 0.32, n = 10) accounted for the remaining heterogeneity in the data.

No relationship was uncovered between fecal GCs and reproductive success with either the standard or phylogenetically informed meta-analyses (Figure 1). Because there were no effect sizes for males, we were unable to test for an effect of sex. Although substantial heterogeneity exists in the data, none of the moderators explained this variation (value of a reproductive bout: β = 0.037, 95% CI = −0.44 – 0.49, *p* = 0.89, n = 12; longevity: β = 0.13, 95% CI = −0.26 – 0.48, *p* = 0.51, n = 12).

### (7) Experimentally elevated GCs and fitness

Both the standard and phylogenetically informed meta-analyses showed that experimentally increasing GCs reduced survival (Figure 1). None of the life history moderators (sex (M): β = −0.25, 95% CI = −0.52 – 0.075, *p* = 0.13, n = 5; value of a reproductive bout: β = 0.46, 95% CI = −0.15 – 0.82, *p* = 0.13, n = 6; longevity: β = −0.0098, 95% CI = −0.023 – 0.003, *p* = 0.13, n =6) explained the remaining heterogeneity in the data; however, this is unsurprising given the small sample size. The effect of GC manipulation on survival was associated with the length of time between administering treatment and assessing survival. More time before assessing survival yielded more negative relationships between GC treatment and survival, though the effect is very small (β = −0.0005, 95% CI = −0.0009 – 0.000, *p* = 0.036, n = 6).

Overall, experimental elevations of GCs reduced reproductive success, but this relationship was not significant after incorporating the expected covariance among related species (Figure 1). None of the life history moderators (sex (M): β = 0.016, 95% CI = −0.45 – 0.47, *p* = 0.95, n = 12; value of a reproductive bout: β = 0.46, 95% CI = −0.85 – 0.98, *p* = 0.58, n = 15; longevity: β = −0.016, 95% CI = −0.049 – 0.017, *p* = 0.34, n = 15) or time from treatment until measuring reproductive success (β = 0.0002, 95% CI = −0.0011 – 0.0016, *p* = 0.73, n = 14) explained the heterogeneity in effect sizes.

### (8) Tests for publication bias

Egger’s test for bias, the trim and fill test for missing data, and an investigation of patterns in effect sizes across time indicate no evidence for publication bias for the meta-analyses of baseline GCs and survival, stress-induced GCs and either survival or reproductive success, fecal GCs and either survival or reproductive success, and GC manipulation and reproductive success (details in the supplementary material).

We observed evidence for time-lag bias in both the overall (Figure 4) and bird-only meta-analyses baseline GCs and reproductive success: in both analyses, effect sizes increased with publication year, generally becoming closer to 0 in more recent years (overall baseline GC-reproductive success: β = 0.03, t = 4.28, *p* < 0.0001; birds-only: β = 0.04, t = 4.39, *p* < 0.0001). Neither Egger’s test nor trim and fill analysis provided evidence for bias for the GC-reproductive success analyses (see supplementary material).

**Figure 4.**
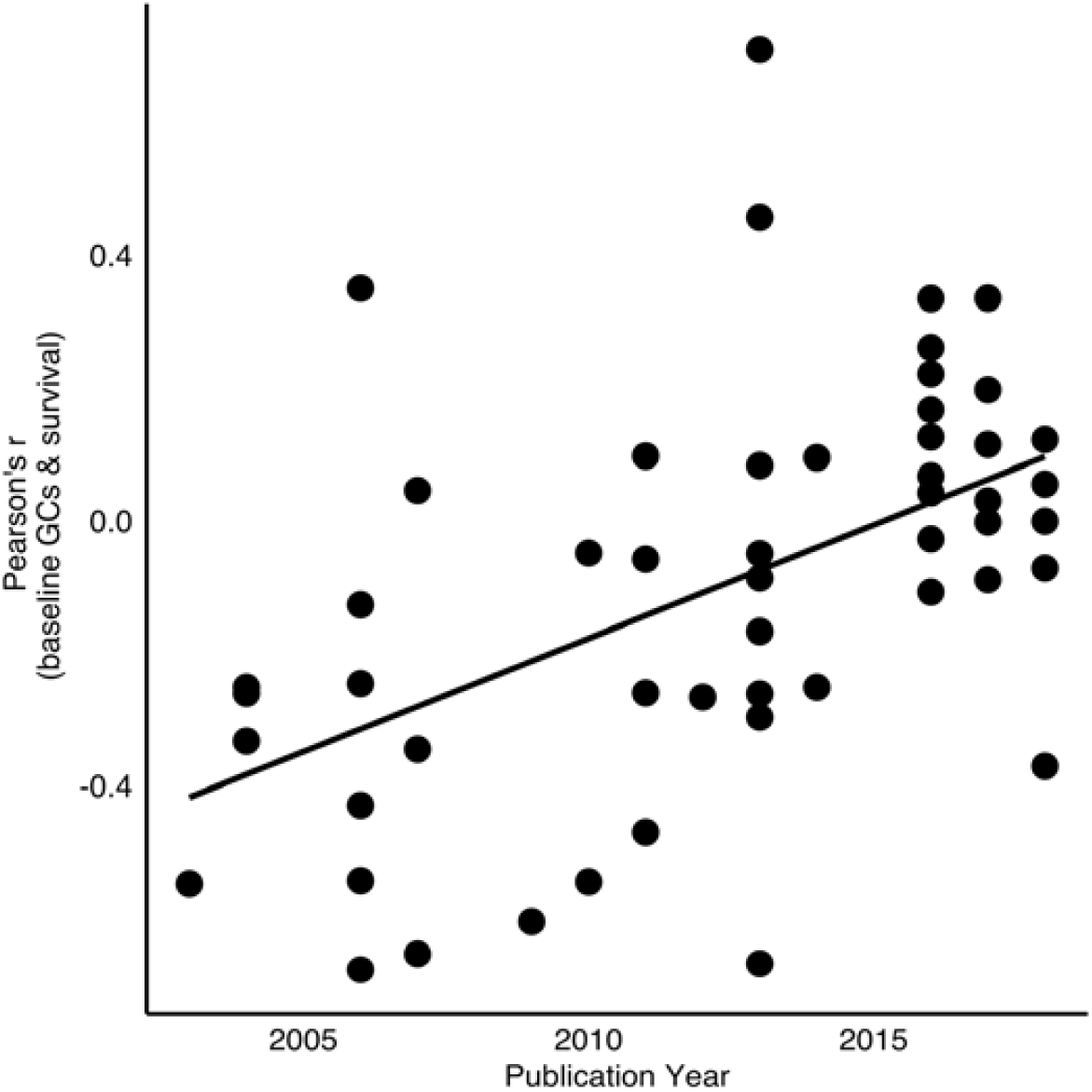
Effect sizes of the relationships between baseline GCs and fitness increase over time, approaching 0. The more negative effect sizes observed in earlier publications could be the result of publication bias favoring more negative GC-fitness relationships.

The test for bias in the meta-analysis of the effects of experimental GC elevation on survival indicated that there is asymmetry in the funnel plot, which can indicate publication bias. Although the Egger’s test provided no evidence of bias for (z = 0.20, *p* = 0.84), the trim and fill analysis suggested that there is some evidence for publication bias and estimated that 1 effect size was missing. The trim-and-fill test allows for “filling” in the missing effect size and recalculating the parameters of the meta-analysis. Adjusting for the missing effect size did not change the overall direction of the relationship between experimentally elevated GCs and survival, but the effect was no longer significant (r = −0.10, 95% CI = −0.26 – 0.07, *p* = 0.24). Because both the effect size and sample size (n = 6) for this analysis were relatively small, they are particularly sensitive to adjusting for missing effect sizes. This result suggests that more data is necessary to draw strong conclusions about the effects of experimental GC elevation on fitness. There was no relationship between publication year and effect sizes (β = −0.013, t = -1.2, *p* = 0.29). No other test of publication bias indicated a need to re-evaluate results, and the details of those analyses are described in the supplementary material.

## IV. Discussion

### (1) Plasma GCs and survival

As predicted, neither baseline nor stress-induced GC levels showed consistent directional relationships with survival. Baseline GC levels tended to negatively predict survival, but this relationship was not significant when incorporating the expected covariance between related species Instead, our results support the prediction that life history will influence the nature of the relationship between GC levels and survival. Among long-lived species, individuals with higher baseline GC levels had lower survival, a pattern that was not apparent in shorter-lived species (Figure 2A). Longer-lived species may suffer greater phenotypic costs of elevated GC levels (including oxidative stress, accelerated telomere shortening, and tissue damage), because these costs take longer to accumulate (Sapolsky, 1999; Metcalfe & Alonso-Alvarez, 2010; Gassen *et al*., 2017). We also predicted that because elevated baseline GC levels may help to support energetically demanding life history stages (Moore & Jessop, 2003; Bonier *et al*., 2009a; Crespi *et al*., 2013), higher baseline GC levels would be relatively more beneficial to reproductive success in species investing more in high value reproductive attempts (Bókony *et al*., 2009; Vitousek *et al*., 2019). Although the data reveal a trend in this direction, the relationship was not significant (Figure 2B). Thus, baseline GC-survival relationships seem to be influenced more strongly by lifespan than by reproductive strategy.

The relationship between stress-induced GCs and survival also differed strongly by life history strategy. Mounting a stronger GC stress response was negatively associated with survival among longer-lived species, but positively associated with survival among shorter-lived species (Figure 3A). This is consistent with the idea that responding effectively to stressors by mounting a strong GC strong response may be relatively more beneficial in organisms facing greater threats to survival (Angelier *et al*., 2009; Hau *et al*., 2010; Schoenle, Zimmer, & Vitousek, 2018; Vitousek *et al*., 2019), as well as the potential for the cumulative costs of elevated GC levels to be greater in longer-lived animals (Lupien *et al*., 2009; Gassen *et al*., 2017). Organisms with higher brood value also showed a more positive relationship between stress-induced GCs and survival than those with lower brood value (Figure 3B). This relationship is not one that we predicted, but is consistent with the idea that GCs mediate trade-offs between investment in survival and reproduction (Wingfield & Romero, 2001; Breuner *et al*., 2008), with a robust GC response conferring a survival benefit among species that invest heavily in a small number of reproductive attempts. As discussed below, high stress-induced GCs appear to negatively impact reproductive success across contexts. Taken together, these results suggest that the GC stress response may be particularly influenced by life history trade-offs in species with high reproductive value (with stronger GC responses favoring survival at a cost to reproduction). In contrast, both survival and reproductive success appear to favor lower GC responders in species that invest less heavily in individual reproductive attempts.

### (2) Plasma GCs and reproductive success

Across studies, individuals with higher GC levels (both baseline and stress-induced) had lower reproductive success. These findings are consistent with the prediction that acute increases in GCs divert resources away from reproduction and towards immediate survival (Wingfield & Sapolsky, 2003; Breuner *et al*., 2008). As discussed above, life history did not influence the relationship between stress-induced or baseline GCs and reproductive success. More surprising was the finding of a consistently negative relationship between baseline GC levels and reproductive success. Although elevated baseline GCs are often seen in individuals facing stressors, baseline GCs are also frequently upregulated in advance of energetically demanding life history stages, and are believed to help organisms mobilize energy to meet these demands (Romero, Reed, & Wingfield, 2000; Bonier *et al*., 2009a; Casagrande *et al*., 2018; Vitousek *et al*., 2019).

In birds, we expected a different relationship between GCs and reproductive success across the incubation and chick-rearing life history stages because of their differing energetic demands (Westerterp & Bryant, 1984; Martin, 1987). Indeed, studies within avian species demonstrate that the relationships between GCs and reproductive success can change across life history stages (Bonier *et al*., 2009b; Ouyang *et al*., 2011; Vitousek *et al*., 2018). Abandoning a reproductive effort during incubation might be less costly than at later stages, particularly if it is possible to quickly initiate another breeding attempt (Love *et al*., 2004). If renesting is possible and the current conditions are unlikely to permit birds to successfully fledge chicks, it might be adaptive for higher baseline GCs during incubation to increase the chances of nest desertion (Lothery *et al*., 2014). Selection might therefore favor a role for high GCs in mediating brood abandonment early in breeding, leading to more negative effects of GCs on reproduction during this period. Alternatively, selection might favor a role for high GCs in mediating successful fledging during chick-rearing because elevated baseline GCs could help to support this energetically demanding period (Bonier *et al*., 2009b). Our analysis found no consistent difference in the relationship between baseline GCs and reproductive success across these life history stages. This could result from variation in the selective pressures facing different species or populations across life history substages. It is also possible that the time-lag bias caused by the publication of substantially more negative effect sizes in baseline GC-fitness relationships in the early 2000s (Figure 4) skewed the results of this analysis, reducing our ability to detect context-dependence.

### (3) Fecal GC-fitness relationships

Observed patterns for fecal GCs and fitnesss can be explained by the expected similarity in trait values of related species; re-analyzing these patterns in a phylogenetic framework eliminated the statistical significance of these patterns. Fecal GCs reflect circulating hormone levels over longer time periods than baseline or peak stress-induced levels. Depending on the context of the organism, longer-term measurements of hormone levels obtained from feces or feathers may be more closely associated with either baseline (Good, Khan, & Lynch, 2003; Cavigelli *et al*., 2005; Bortolotti *et al*., 2008) or stress-induced GCs (Harper & Austad, 2002; Fairhurst *et al*., 2013). Although the relationships between GCs and survival are complex and context-dependent, we saw a consistently negative relationship between reproductive success and both baseline and stress-induced GCs. Why then do we not see this relationship in fecal GCs? One possibility is that the presence of these relationships is obscured by inter-individual variation in diet, metabolic rate, or hormone metabolite formation (Goymann, 2012). Measured fecal GC concentrations can also be affected by variation in the time between defecation and collection/freezing, and by environmental factors such as temperature, precipitation, and microbial content (Dantzer *et al*., 2014).

### (4) Experimentally elevated GCs and fitness

Experimentally elevated GC levels were consistently negatively associated with survival. Individuals with manipulated GC levels tended to have lower reproductive success – a pattern that mirrors the relationships seen in unmanipulated individuals – but this effect was not significant in analyses that incorporated phylogenetic relationships. Experimentally altering GC levels consistently reduced survival, a pattern that is interesting considering that natural variation in GCs showed variable and context-dependent relationships with survival. This discrepancy could result from several factors. Hormone manipulations often do not accurately mimic natural variation in circulating GCs across various temporal scales (Fusani, 2008). Additionally, because hormone regulation can be adaptively modulated in accordance with the internal and external context of an individual, it is not surprising that manipulations that move individuals away from their natural regulatory patterns could negatively impact fitness, even if similar relationships are not seen in unmanipulated individuals (Crossin *et al*., 2016; Schoenle *et al*., 2019). Intriguingly, experimentally altered GC levels were associated with a greater decrease in survival in studies that measured survival over longer time periods. This is consistent with the potential for many of the salient costs of GCs to accumulate over longer timescales. Although most studies assess only the short-term survival impacts of GCs, these patterns suggest that GC manipulations could be more costly than previously assumed.

## V. CONCLUSIONS

Our results suggest strong context-dependence in glucocorticoid-survival relationships, and more consistently negative relationships between plasma GCs and reproductive success. Unlike measurements of plasma GCs, fecal GCs did not show consistent relationships with either metric of fitness. Our analyses also support the potential for GC elevation to have pathological effects: experimental GC elevation consistently reduced survival, whereas its effect on reproductive success was less clear.

Life history context appears to be crucial to understanding the relationships between GCs and fitness, but as predicted, the role of life history varies across fitness metrics. Although plasma GCs were not consistently associated with survival, the direction of GC-survival relationships were strongly dependent on life-history. In contrast, plasma GCs were negatively associated with reproductive success, but life history context was not associated with the variation in the direction of the relationship. We recommend that moving forward, researchers aim to explicitly test how the optimal GC response to challenges varies across contexts, including those not measured in these meta-analyses. Studies manipulating stressors (including the frequency and intensity of these stressors) and GCs across environmental gradients as well as life history stages and strategies could provide insight into how different contexts may influence GC–fitness relationships.

Finally, our search of the literature revealed gaps in our knowledge of GC-fitness relationships. Nearly all observational studies were conducted during the breeding season, and more than twice as many of the available effect sizes were from tests of the GC-reproductive success relationship than of the GC-survival relationship (Table 1). Furthermore, the available GC-fitness effect sizes are highly biased towards birds, even though it seems like relevant data should exist for other taxa (Beehner & Bergman, 2017). To develop a more comprehensive understanding of the relationships between GCs and fitness, we also encourage future studies to extend beyond the breeding season and to incorporate survival analyses in more diverse taxa.

## Supporting information

Supplementary Methods and Results

data set

## Acknowledgments

We thank Dan Becker for providing advice on methods for meta-analyses. We thank the HormoneBase Consortium and Creagh Breuner for discussions on this topic.

