## Supplementary Methods and Results for "Does variation in glucocorticoid regulation predict fitness? A phylogenetic meta-analysis"

**Supplemental Methods**

**Tests for significance of random effects**

For each meta-analysis, we tested the importance of the reference ID, species, and whenever relevant, group ID using likelihood ratio tests. We separately tested each random variable for significance within an intercept-only meta-analysis. Variables that were found to be significant in these individual analyses were then included in an intercept-only model. We checked which of these random effects remained significant after accounting for the others, and only retained those random effects in the general models.

*Baseline GCs and survival*

Neither reference ID (*χ*^2^ = 2.39, *p* = 0.12), group ID (*χ*^2^ = 1.51, *p* = 0.22), nor species (*χ*^2^ = 2.68, *p* = 0.10) significantly improved model fit when entered individually as random effects, and they were not included in the meta-analyses.

*Baseline GCs and reproductive success*

Group ID improved model fit when entered individually into the model (*χ*^2^ = 54.45, *p* < 0.0001), but reference ID and species did not have significant effects above the effect of Group ID when they were entered into the models simultaneously (reference ID: *χ*^2^ = 0, *p* = 1; species: *χ*^2^ = 0, *p* = 1). Thus, group ID was retained in all analyses for baseline GCs and reproduction.

*Baseline GCs and reproductive success – birds only*

Group ID improved model fit (*χ*^2^ = 44.77, *p* < 0.0001), but reference ID and species did not have significant effects above Group ID when they were entered into the models simultaneously (reference ID: *χ*^2^ = 0, *p* = 1; species: *χ*^2^ = 0, *p* = 1). Thus, group ID was retained as a random effect for the meta-analyses.

*Stress-induced GCs and survival*

Neither reference ID (*χ*^2^ = 0.21, *p* = 0.65), group ID (*χ*^2^ = 2.33, *p* = 0.13), nor species (*χ*^2^ = 0.21, *p* = 0.65) significantly improved model fit when entered individually as random effects, and thus, they were not included in the meta-analyses.

*Stress-induced GCs and reproductive success*

Neither reference ID (*χ*^2^ = 0.000, *p* = 1), group ID (*χ*^2^ = 0.53, *p* = 0.47), nor species (*χ*^2^ = 0.000, *p* = 1) significantly improved model fit when entered individually as random effects, and they were not included in the meta-analyses.

*Fecal GCs and survival*

Reference ID accounted for differences in group ID and species. Thus, only reference ID was used as a random effect; it significantly improved the model (*χ*^2^ = 66.78, *p* < 0.001) and was retained for the analyses.

*Integrated GCs and reproductive success*

Reference ID accounted for differences in group ID and species and thus, only reference ID was tested as a random effect. Reference ID significantly improved the model (*χ*^2^ = 22.68, *p* < 0.001) and was included in the analyses.

*Experimentally elevated GCs and survival*

Neither reference ID (*χ*^2^ = 0, *p* = 1), group ID (*χ*^2^ = 0, *p* = 1), nor species (*χ*^2^ = 0, *p* = 1) significantly improved model fit when entered as random effects and were not included in the meta-analyses.

*Experimentally elevated GCs and reproductive success*

Reference ID (*χ*^2^ = 6.25, *p* = 0.01), but not group ID (*χ*^2^ = 1.45, *p* = 0.23) or species ID (*χ*^2^ = 3.71, *p* = 0.054) significantly improved model, and thus reference ID was included as a random effect.

**Supplemental Results**

**Tests for publication bias**

Table S1: Results of three publication bias tests including Egger’s Test, Trim and Fill Analyses, and linear regression of publication year and effect size. Tests indicating possible publication bias are in bold and are explained in detail in the main text.

| **Meta-analysis** | **Egger's Test** | | **Trim & Fill Analysis** | | **Regression of Effect Sizes and Publication Year** | | |
| --- | --- | --- | --- | --- | --- | --- | --- |
|  | ***z*** | ***p*** | **No. Missing Effect Sizes** | **Adjusted Effect Size** | **β** | **t** | ***p*** |
| Baseline GCs & Survival | -0.64 | 0.53 | 0 | NA | 0.02 | 2.09 | 0.06 |
| Baseline GCs & Reproduction | -0.38 | 0.7 | 0 | NA | **0.03** | **4.28** | **<0.0001** |
| Baseline GCs & Reproduction (birds only) | -0.4 | 0.69 | 0 | NA | **0.04** | **4.39** | **<0.0001** |
| Stress-induced GCs & Survival | 0.7 | 0.48 | 3 | r = 0.04, 95% CI = -0.07 – 0.14, p = 0.48 | -0.0004 | -0.22 | 0.83 |
| Stress-induced & Reproduction | 1.56 | 0.12 | 5 | r = -0.16, 95% CI = -0.26 – -0.06, p = 0.001 | -0.0068 | -0.54 | 0.6 |
| Fecal GCs & Survival | 0.48 | 0.63 | 2 | r = -0.31, 95% CI = -0.51 – -0.10, *p* = 0.004 | -0.00015 | -0.007 | 0.99 |
| Fecal GCs & Reproduction | -0.49 | 62 | 2 | r = -0.017, 95% CI = -0.12 – 0.08, *p* = 0.74 | -0.0006 | -0.048 | 0.96 |
| Experimentally Elevated GCs & Survival | 0.2 | 0.84 | **1** | **r = -0.10, 95% CI = -0.26 – 0.07, *p* = 0.24** | -0.013 | -1.2 | 0.29 |
| Experimentally Elevated GCs & Reproduction | -1.14 | 0.25 | 0 | NA | 0.019 | 0.8 | 0.44 |
